# A sensitive fluorometric assay to detect aldo-keto reductase and carbonyl reductase activity based on a naphthaldehyde derivative

**DOI:** 10.64898/2026.06.15.732224

**Authors:** Lucia Piazza, Raquel Pequerul, Xavier Parés, Francesco Balestri, Giovanni Signore, Antonella Del Corso, Jaume Farrés

## Abstract

We have developed a fluorometric assay for detecting reductase activity in biological samples through 4-methoxy-1-naphthalenemethanol (MONOL-41) formation. The enzyme carbonyl reductase 1 (CBR1) and four members of the aldo-keto reductase (AKR) 1 family (AKR1A1, AKR1B1, AKR1B10, AKR1C3) were evaluated for their ability to reduce 4-methoxy-1-naphthaldehyde (MONAL-41). AKR1B1 and CBR1 followed Michaelis–Menten kinetics, whereas AKR1B10, AKR1A1, and AKR1C3 showed substrate inhibition above 10 µM (70 µM for AKR1C3). Among the tested enzymes, AKR1B10 displayed the highest catalytic efficiency in the absence of substrate inhibition. The MONOL-41 assay was compared with the standard NADPH-based method, showing improved sensitivity, robustness, and lower detection limits (0.77 µg/mL *vs.* 1.49 µg/mL). These results confirm its suitability for monitoring AKR1B10 activity. The assay was then applied to A549 cell extracts, which express multiple reductases. Activity decreased at substrate concentrations above 10 µM, suggesting a predominant role of AKR1B10. Inhibition studies using tolrestat and high MONAL-41 concentrations indicated a limited contribution of CBR1 (∼7–8%). Considering both catalytic efficiency and expression levels, AKR1B10 appears to be the main contributor to reductase activity in this model. In A549 living cells, MONAL-41 showed no cytotoxicity up to 50 µM and enabled real-time monitoring due to its membrane permeability. However, oxidation by aldehyde dehydrogenases can generate MONOIC-41, which has similar spectral properties but a lower quantum yield, potentially affecting signal interpretation. Overall, this assay represents a sensitive and cost-effective tool for detecting reductase activity and screening inhibitors.

## 1. Introduction

The conversion of endogenous carbonyl compounds and xenobiotics to their corresponding alcohols is primarily catalyzed by NADPH-dependent enzymes belonging to aldo-keto reductase (AKR) and short chain dehydrogenase/reductase (SDR) superfamilies [1, 2]. Carbonyl-reducing enzymes play crucial roles in several processes such as carcinogenesis, drug resistance, inflammation, signal transduction, and detoxification of toxic aldehydes [3–5]. Aldo-keto reductase 1 family members (AKR1A1, AKR1B1, AKR1B10 and AKR1C3) and carbonyl reductase 1 (CBR1) are involved in the detoxification of reactive aldehydes and ketones, thereby contributing to cellular protection against oxidative stress. AKRs represent a functionally diverse group with distinct substrate specificities, participating in processes such as glucose metabolism, lipid peroxidation-derived aldehydes detoxification, drug metabolism and steroid hormone regulation [1, 6]. In contrast, CBR1 belongs to the short-chain dehydrogenase/reductase (SDR) family and is broadly implicated in drug metabolism, including anticancer drugs such as anthracyclines [7]. As a result, their involvement in cellular pathways and their potential as drug targets are actively being studied [8–10].

Standard procedures used to perform the kinetic characterization of reductases are based on monitoring the rate of NADPH oxidation in the presence of carbonyl-bearing substrates [11]. Although these methods provide accurate measurements for purified preparations, the lack of specific probes could hamper the detection of a targeted enzymatic activity in biological samples where multiple NADPH-dependent reductases with shared substrate specificities are present. Therefore, a probe capable at discriminating the activity of specific reductases in cell lysates is needed.

Wierzchowski *et al.* [12] developed a fluorometric method with a demonstrated specificity for the detection of the reductase activity of ADH class I isoforms in human serum and the liver of rats poisoned with methanol [13, 14]. The assay was based on the peculiar spectral properties of the fluorogenic naphthalene derivative 4-methoxy-1-naphthaldehyde (MONAL-41) and its corresponding alcohol 4-methoxy-1-naphthalenemethanol (MONOL-41). MONAL-41 is poorly fluorescent (quantum yield, Φ = 0.003), but its reduction to MONOL-41 results in a marked spectral change with a fluorescent quantum yield (Φ = 0.36) significantly superior to several fluorescent or fluorogenic probes, including NAD(P)H [12]. Additionally, MONOL-41 can be effectively excited at λ>310 nm, preventing a significant masking effect by proteins contained in biological samples [12]. Accordingly, the sensitivity of the assay in detecting class I alcohol dehydrogenase (ADH) isoforms in biological samples has been demonstrated in multiple studies and it is significantly superior to assays based on NAD(P)H absorbance and fluorescence [13–15]. However, to the best of our knowledge, the potential of MONAL-41 as a probe to detect the CBR1 or AKR1 activity in biological samples has not yet been explored. Naphthaldehydes and other polycyclic aromatic compounds have been reported previously to be substrates of AKR1 isoforms and CBR1 [16–18], although detailed kinetic analysis with MONAL-41 is lacking. In contrast to the standard procedures, the assay based on monitoring MONAL-41 reduction is promising in terms of sensitivity and may allow the detection of specific reductases in biological samples. Moreover, MONAL-41 shows an optimal predicted membrane permeability (XlogP3 = 3.1 [19]), suggesting the additional advantage to perform live-cell detection of reductases activity. Our study aims to refine an innovative fluorometric procedure to detect the activity of reductases with a focus on the specificity and the optimization of sensitivity, repeatability and cost/time-effectiveness.

## 2. Materials and methods

### 2.1. Materials

NADPH tetrasodium salt was purchased from Biosynth (Nobelova, Bratislava, Slovakia); D,L-dithiothreitol (DTT) was purchased from MP Biomedicals, LLC (Illkirch, France); isopropyl-β-D-1-thiogalactopyranoside (IPTG) and phenylmethylsulfonyl fluoride (PMSF) were obtained from Merck Life Science (Milan, Italy); D,L-glyceraldehyde, bovine serum albumin (BSA), dimethyl sulfoxide (DMSO) and YM10 ultrafiltration membranes were purchased from Merck-Millipore (Darmstadt, Germany); 9,10-phenanthrenequinone, 4-nitrobenzaldehyde, *trans*-2-nonenal, were purchased from Sigma chemical company (St Louis, MO, USA); glutathionyl (GS)-nonanal was synthesized as previously described [20]; 4-methoxy-1-naphthaldehyde (MONAL-41) and 4-methoxy-1-naphthalenemethanol (MONOL-41) were purchased from CymitQuimica (Barcelona, Spain). Presto Blue ™ Cell Viability Reagent was purchased from Invitrogen (Carlsbad, CA, USA). Sodium dihydrogen phosphate, acetonitrile, and ethanol were purchased from Carlo Erba Reagents (Milan, Italy). 96-well F-bottom black plates were purchased from Greiner Bio-One (Frickenhausen, Germany). The A549, a human lung adenocarcinoma epithelial cell line, was from ATCC (Rockville, MD, USA). All other chemicals were of reagent grade.

### 2.2. Purification of human recombinant AKRs and CBR1

The human recombinant AKR1B1 and CBR1 were expressed and purified to electrophoretic homogeneity, as previously described [8, 20, 21]. The gene sequences of human AKR1A1, AKR1B10 and AKR1C3 were individually cloned into pET30 plasmids by Eurofins Genomics (Ebersberg, Germany) under the control of an inducible promoter activated by IPTG. The constructs also contained a gene conferring resistance to kanamycin. The BL21 strain of *E. coli* competent cells (Agilent, Santa Clara, CA, USA), was used for transformation with each plasmid. BL21 cells were incubated with the pET-30 vector for 1 h on an ice bath, followed by heat shock treatment for 60 s at 42°C and immediate transfer to an ice bath for 5 min. Then, 10 volumes of Luria-Bertani (LB) broth (10 g/L tryptone, 5 g/L yeast extract and 10 g/L NaCl) were added to the cells, which were further incubated at 37°C for 1 h in agitation. Transformed cells were grown overnight at 37°C in LB broth supplemented with 30 μg/mL kanamycin. When the culture had reached an O.D.600 of 0.8, recombinant enzyme expression was induced by addition of 0.4 mM IPTG, and the culture was kept for 3 h at 37°C. Then, cells were centrifuged at 3,500 *xg* for 15 min at 4°C. The resulting pellet was resuspended (1 g/4 mL) in 20 mM Tris-HCl buffer, pH 7.4, containing 2 mM DTT and 1 mM PMSF. The resulting suspension after three freezing (−20°C for 30 min) and thawing (37°C for 90 s) cycles was subjected to five cycles of sonication (10 s with 20 s intervals). The homogenates were finally centrifuged at 10,000 *xg* for 90 min at 4°C and the obtained supernatants were referred to as the crude extracts.

The different crude extracts (approximately 85 mg of protein each), containing the proper recombinant enzyme, were subjected to a purification procedure consisting of two sequential chromatographic steps (both performed at 4°C). In the first step the extract was loaded on a DEAE Sepharose (Merck Life Sciences) column (3.5 x 15 cm), previously equilibrated with 50 mM sodium phosphate buffer, pH 7.0 containing 2 mM DTT. The elution was performed with the same buffer at a flow rate of 20 mL/h. Fractions of 3 mL were collected, and absorbance at 280 nm was monitored. Each fraction was assayed for enzymatic activity using the appropriate enzymatic assay (detailed in Section 2.3). Fractions displaying activity were pooled and concentrated using a YM10 Amicon ultrafiltration membrane and then applied to a Blue Sepharose column (Merck Life Science) (1.6 x 6.5 cm,) pre-equilibrated with 50 mM sodium phosphate buffer, pH 7.0, at a flow rate of 20 mL/h. Fractions of 2 mL were collected and absorbance at 280 nm was monitored. Each fraction was assayed for enzymatic activity using the appropriate enzymatic assay. When the absorbance at 280 nm reached baseline levels, a 50 mM sodium phosphate buffer, pH 7.0, containing 0.37 M NaCl was applied. When the absorbance reached again baseline levels, the recombinant enzyme was eluted with 50 mM sodium phosphate buffer, pH 7.0, containing 2 mM DTT, 0.37 M and 0.1 mM NADPH. Fractions containing enzymatic activity were pooled and concentrated using a YM10 ultrafiltration membrane. Purity assessment was performed through SDS-PAGE analysis followed by Coomassie Brilliant Blue staining. Before use, the purified preparations were extensively dialyzed against 10 mM sodium phosphate buffer, pH 7.0. In the case of AKR1B10 and AKR1C3, the dialysis buffer was supplemented with 0.5 mM DTT.

### 2.3. Standard enzymatic assay

The standard assays of recombinant enzymes were performed at 37°C, monitoring the decrease in absorbance at 340 nm linked to NADPH oxidation (ε340 = 6.22 mM^−1^·cm^−-1^) by a Biochrom Libra S60 UV-visible spectrophotometer. The composition of the standard assay mixtures is reported in Table 1.

**Table 1.** Standard assay mixtures of the human recombinant AKR1A1, AKR1B1, AKR1B10, AKR1C3 and CBR1.

| Enzyme | Composition of standard assay mixtures |
| --- | --- |
| AKR1A1 | 100 mM sodium phosphate buffer, pH 7.0, 0.18 mM NADPH, and 0.7 mM 4-nitrobenzaldehyde |
| AKR1B1 | 250 mM sodium phosphate buffer, pH 6.8, 0.18 mM NADPH, 0.4 M ammonium sulphate, 0.5 mM EDTA, and 4.7 mM D,L-glyceraldehyde |
| AKR1B10 | 100 mM sodium phosphate buffer, pH 7.0, 0.18 mM NADPH, and 15 mM D,L-glyceraldehyde |
| AKR1C3 | 50 mM sodium phosphate buffer, pH 6.0, 0.18 mM NADPH, 12 μM 9,10-phenanthrenequinone |
| CBR1 | 50 mM sodium phosphate buffer, pH 6.0, 0.18 mM NADPH, 30 μM GS-nonanal |

The concentration of the active protein was determined with the following equation:

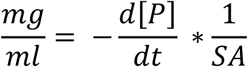

Where 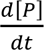 is the rate of change of the NADPH oxidation (µmol · mL^−1^ · min^−1^) and SA is the specific activity (µmol · min^−1^ · mg^−1^) of the recombinant enzyme for the substrate used in the standard assay. The specific activities of the recombinant enzymes are reported in Table 2.

**Table 2.** Specific activity of the human recombinant AKR1A1, AKR1B1, AKR1B10, AKR1C3 and CBR1 for the substrates used in their respective standard assay.

| Enzyme | Specific activity<br>($\mu\text{mol} \cdot \text{min}^{-1} \cdot \text{mg}^{-1}$) |
| --- | --- |
| AKR1A1 | 27.4 |
| AKR1B1 | 6 |
| AKR1B10 | 1.6 |
| AKR1C3 | 0.41 |
| CBR1 | 35 |

### 2.4. Preparation of fluorogenic substrate and internal standard solution

MONAL-41 stock solutions (3-6 mM) were prepared in a water:acetonitrile mixture (2:1 v/v). The internal standard MONOL-41 stock solutions (1.8 mM) were prepared in ethanol. The molar extinction coefficients of MONOL-41 and MONAL-41 (ε296= 7.1 mM^−^1·cm^−1^ and ε333= 12.81 mM^−^1·cm^−1^, respectively) were determined in the respective stock solvents by a Biochrom Libra S60 spectrophotometer.

### 2.5. Cell culture and extract preparation

A549 cell line was grown at 37°C under a humidified atmosphere of 5% CO2 and 95% air in DMEM High Glucose medium, supplemented with 10% FBS and 2 mM glutamine. Cells were washed with PBS containing 1 mM PMSF before being collected and stored at −80°C. Cell extracts were obtained through three cycles of freezing and thawing and then centrifuged at 16000*xg* for 15 min at 4°C to remove cell debris. Prior to use, cell extracts were extensively dialyzed against 10 mM sodium phosphate buffer, pH 7.0.

### 2.6. MONOL-41 fluorometric assay of recombinant enzymes and cell extracts

The fluorometric assay of both purified enzymes and cell extracts were conducted using the FLUOstar Omega (BMG Labtech GmbH) plate reader in 96-well flat-bottom black Greiner microplates and optimized as described in Supplementary Materials. The assay mixtures contained 0.1 M sodium phosphate buffer, pH 7.0, 0.08 mM NADPH, and MONAL-41, in a total volume of 100 μL. The MONAL-41 stock solution was appropriately diluted in the assay buffer before being added to the assay mixtures maintaining acetonitrile concentration below 1% (v/v), which did not affect significatively the enzyme activity. The fluorescence emission linked to the production of MONOL-41 was monitored at 410 nm (bandwidth: 10 nm) by exciting at 325 nm (bandwidth: 40 nm) nm at 37°C. The absolute reaction rate was calculated using the equation:

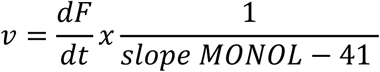

Where 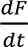 represents the net time-dependent fluorescence slope (i.e., the value obtained by the mix containing the enzyme subtracted of the value obtained from the corresponding reference mix without enzyme), and “slope MONOL-41” is the slope of the calibration curve of MONOL-41 fluorescence in 0.1 M sodium phosphate buffer, pH 7.0, supplemented with 0.08 mM NADPH.

### 2.7. Evaluation of cell viability

The cytotoxicity of the substrate MONAL-41 and its solvent, acetonitrile, on the A549 cell line, was determined by the PrestoBlue™ assay. Five thousand cells per well were seeded on a 96-well plate in a final volume of 100 μL of High Glucose DMEM medium + GlutaMAx supplemented with 10% (v/v) ultracentrifuged FBS and incubated overnight at 37°C and 5% CO2 in air. Various concentrations of acetonitrile and MONAL-41 were prepared in a complete medium. After overnight incubation, the medium was removed and 100 μL of the corresponding dilution was added to cells. Each concentration was tested in triplicate. Cells were incubated for 24 h at 37°C and 5% CO2 in air. After incubation,10 μL of PrestoBlue™ reagent was added to each well and incubated for 3 h. Then, the fluorescence product generated by cells was read in a Tecan Spark Multiplate Reader, with excitation at 531 nm and emission at 572 nm. Viability was determined as the percentage of an untreated control.

### 2.8. MONOL-41 fluorometric assay in intact cells

Reduction of MONAL-41 in cells was determined by monitoring fluorescence emission in conditioned medium. 800,000 cells were seeded on a Petri dish in a final volume of 10 mL of High Glucose DMEM + GlutaMAX supplemented with 10% (v/v) ultracentrifuged FBS and incubated at 37°C and 5% CO2 in air. After overnight incubation, the medium was replaced with 10 mL of complete medium containing 50 μM MONAL-41 dissolved in acetonitrile at a final concentration of 0.53% (v/v) in the medium. Control samples were prepared with the same final solvent percentage without addition of MONAL-41. Detection of MONOL-41 was performed by exciting at 296 nm (path length = 0.3 cm) the conditioned medium collected at several time points from cell cultures treated in both conditions. Fluorescence emission spectra were recorded in the 310-500 nm interval. To correct the fluorescence emission spectra of medium obtained from cell cultures treated in both conditions, the fluorescence emission of medium obtained from two Petri dishes without cells treated with complete medium containing 50 μM MONAL-41 or 0.53% (v/v) of acetonitrile was subtracted as a blank.

### 2.9. Kinetic analysis

The kinetic parameters *kcat* and *KM* were evaluated by non-linear regression analysis of rate measurements *vs.* substrate concentration according to Michaelis-Menten equation through GraphPad Prism 8 software. For enzymes affected by substrate inhibition, the kinetic parameters were obtained by fitting the experimental points at which no inhibitory effect was evidenced.

### 2.10. Other methods

Protein concentration of cell extracts was determined by the Bradford method [22], using BSA as a standard protein.

## 3. Results and discussion

### 3.1. Kinetic properties of MONAL-41 as a substrate of AKR1 and CBR1

Our study aimed to implement an innovative fluorometric assay to detect the activity of reductases in biological samples, through the measurement of the produced MONOL-41. In this regard, we selected a member of SDR superfamily, CBR1, and four members belonging to the aldo-keto reductase family 1 (AKR1), namely A1, B1, B10, and C3 as representative enzymes to test their ability to reduce MONAL-41. The reaction rates of these enzymes reported as a function of MONAL-41 concentration showed different trends (Figure 1): AKR1B1 and CBR1 displayed classical Michaelis-Menten kinetics. On the other hand, substrate inhibition for AKRB10, AKR1A1 and AKR1C3 was observed for MONAL-41 concentration higher than 10 µM for the first two and 70 µM for the latter. Fitting the data in the concentration range where no inhibitory effect was observed allowed the identification of AKR1B10 as the most efficient biocatalyst among the reductases tested (Table 3), being 3.8-fold more efficient than AKR1A1 and about two orders of magnitude more efficient compared with the other tested enzymes.

**Figure 1.**
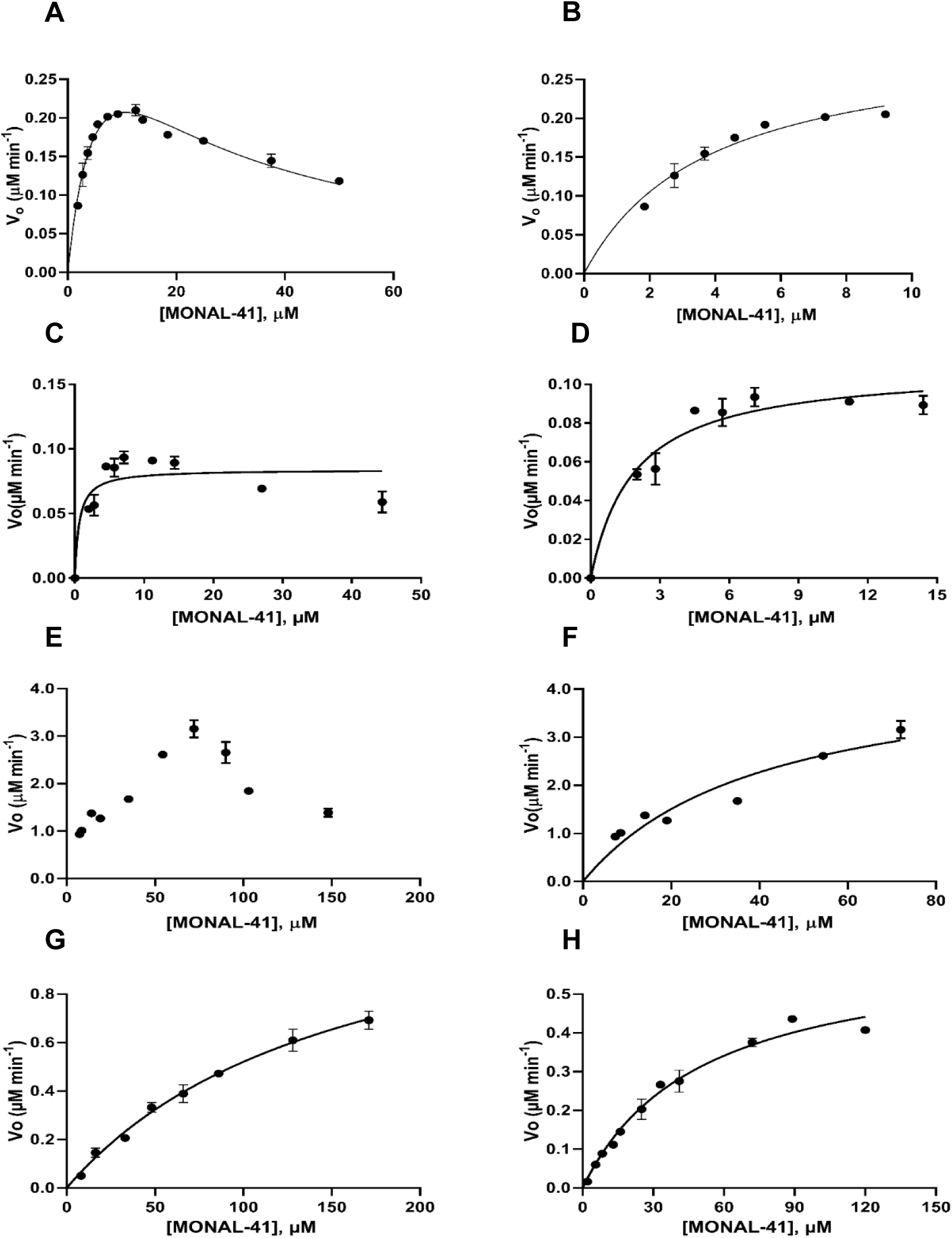
Graphical representation of enzyme activity rate against MONAL-41 concentration. AKR1B10 (panel A), AKR1A1 (panel C), AKR1C3 (panel E), CBR1 (panel G), and AKR1B1 (panel H). Panels B, D, and F show the fit of the experimental points for AKR1B10, AKR1A1 and AKR1C3, respectively, at which no inhibitory effect was evidenced. Results are reported as the mean ± standard deviation of at least three measurements.

**Table 3.** Kinetic parameters of enzymes tested using MONAL-41 as a substrate.

| Enzyme | $K_M$ ($\mu\text{M}$) | $k_{cat}$ ($\text{min}^{-1}$) | $k_{cat}/K_M$ ($\mu\text{M}^{-1} \cdot \text{min}^{-1}$) |
| --- | --- | --- | --- |
| **AKR1B10** | $3.81 \pm 0.50$ | $14.5 \pm 0.8$ | $3.8 \pm 0.7$ |
| **AKR1A1** | $1.49 \pm 0.48$ | $1.56 \pm 0.10$ | $1.0 \pm 0.4$ |
| **AKR1B1** | $50.0 \pm 4.7$ | $2.72 \pm 0.01$ | $0.054 \pm 0.005$ |
| **AKR1C3** | $41.18 \pm 9.55$ | $0.54 \pm 0.06$ | $0.013 \pm 0.005$ |
| **CBR1** | $156 \pm 20$ | $14 \pm 1.0$ | $0.09 \pm 0.02$ |

Kinetic constants were determined using 0.1 M sodium phosphate buffer, pH 7.0, 0.08 mM NADPH, at 37°C, and expressed as mean ± standard deviation of at least three measurements.

### 3.2. Improved sensitivity of the MONOL-41 assay

The sensitivity of the MONOL-41 assay in detecting AKR1B10 activity was tested against that of an assay using the substrate commonly used in the standard assay, following the decrease in fluorescence linked to NADPH oxidation. To this end, AKR1B10 concentrations ranging from 0.2 to 1.13 µg/mL were tested with either the highest non-inhibitory concentration of MONAL-41 (10 µM) or a saturating concentration (15 mM) of glyceraldehyde (Figure 2). The higher brightness of MONOL-41, which is approximately 15-20 times that of NAD(P)H, suggested that the MONOL-41-assay could be more sensitive compared with the standard cofactor-based assay, with the additional advantage of a more selective substrate. When tested with different concentrations of AKR1B10, MONOL-41 showed-sensible improvements in robustness of fit compared with the standard assay (Figure 2).

**Figure 2.**
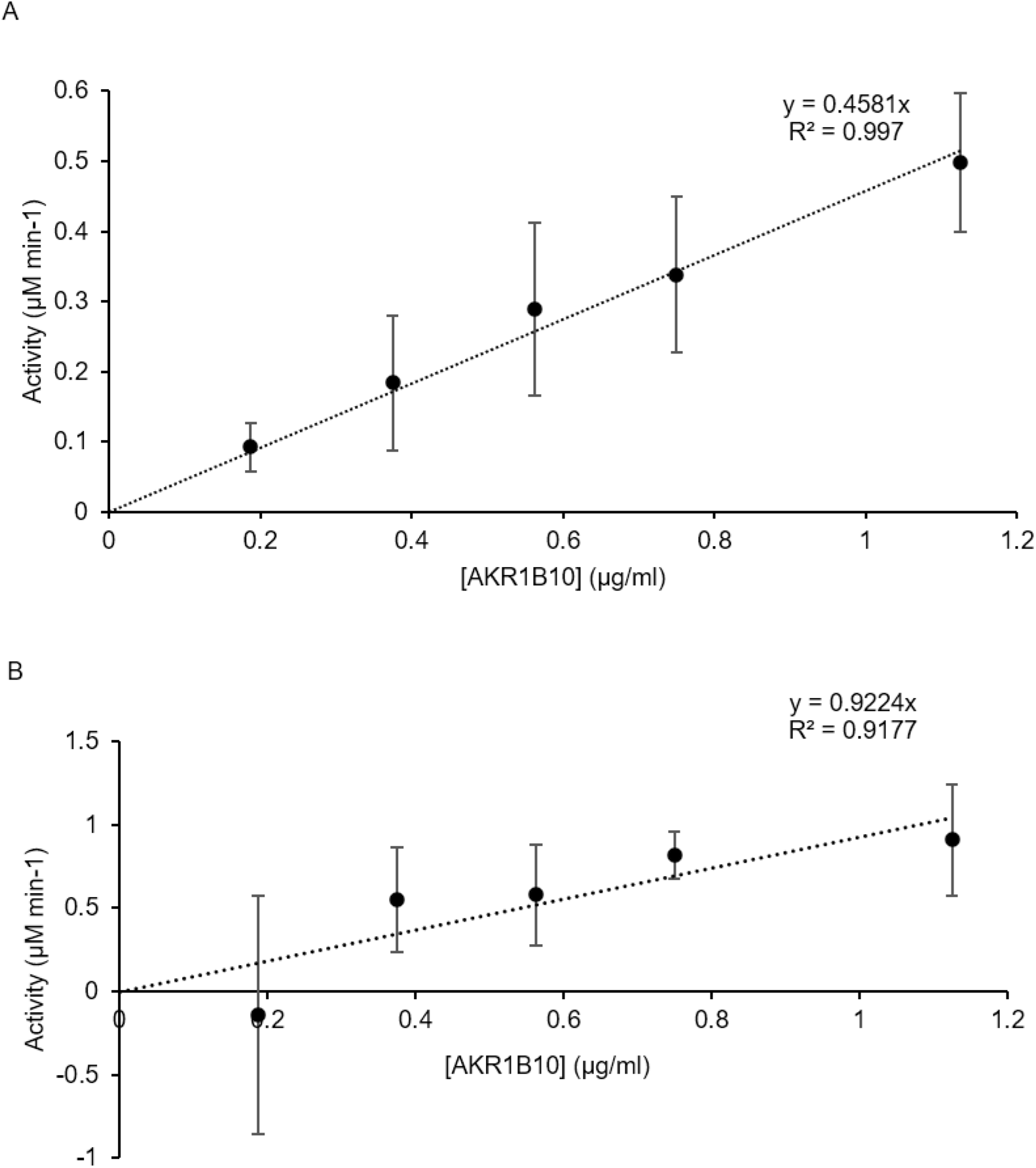
Graphical representation of enzyme activity rate against AKR1B10 concentration. 10 µM MONAL-41 (Panel A) or 15 mM glyceraldehyde (Panel B) was used as a substrate. Same conditions were adopted for both methods, except for the set-up of excitation and emission wavelengths which was optimized for detecting MONOL-41 (325-40 nm and 410-10 nm, respectively) or NADPH (340-10 nm and 460 nm, respectively). Calibration curves of standard MONOL-41 and NADPH are reported in Figure S4. Values are expressed as the mean ± standard deviation of at least three measurements.

A signal-to-noise ratio value of 3.3 is a good estimation of the limit of detection [23], which is reached at concentrations of approximately 70 µg/mL with the MONOL-41 assay. In contrast, the standard assay performed barely reaches a signal-to-noise ratio value of 2 at 1.13 µg/mL (Figure 3).

**Figure 3.**
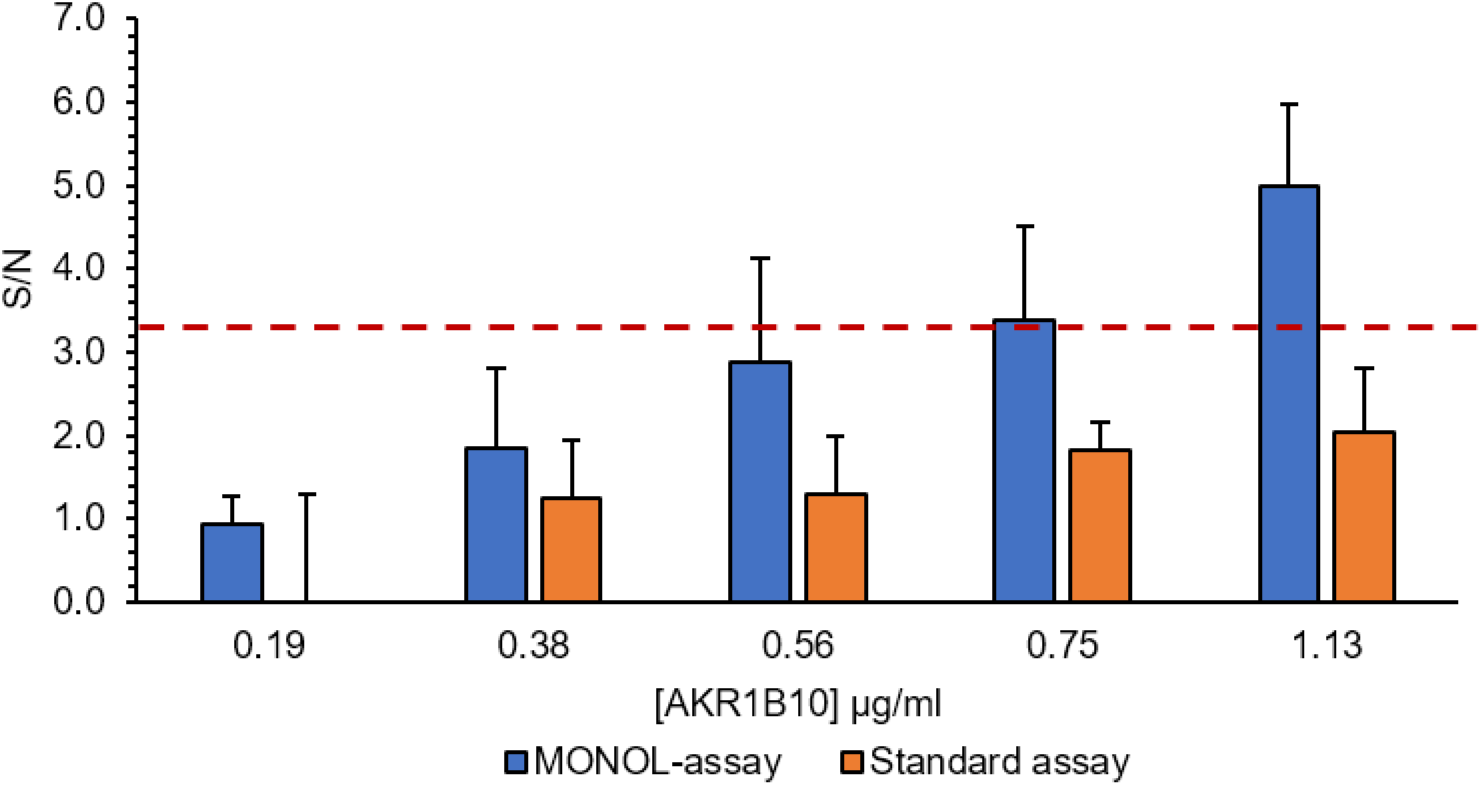
Comparison of the sensitivity between the MONOL-41-assay and the standard NADPH-based method. MONOL-41 assay (blue bars) and NADPH assay (orange bars). Two-way ANOVA test analysis of Signal-to-Noise ratio (S/N) was obtained by the following equation 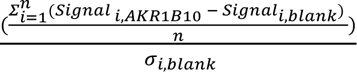, where *Signal*_*i,AKR*1*B*10_ is the RFU · min ^−1^ evaluated for the assay mix containing AKR1B10, *Signal*_*i,blank*_ is the RFU· min*^−^*^1^ evaluated for the assay mix without the enzyme, and *σ*_*i,blank*_ is the standard deviation evaluated for the *Signal*_*blank*_. The red line represents the cut-off for estimating the limit of detection, as reported [23]. The results are reported as the mean ± standard deviation of four independent measurements.

Accordingly, the limits of detection calculated as 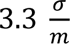 (where σ is the standard deviation of the blank signal and *m* is the slope of the calibration curve of the analyte, according to [23]) are 0.77 µg/mL and 1.49 µg/mL for the MONOL-41 assay and the standard assay, respectively. In our hands, this assay clearly demonstrated the suitability of MONAL-41 as a substrate for AKR1B10 reductase activity, with a sensitivity that outperformed the standard assay.

### 3.3. Detection of reductase activity in A549 cell extracts

Attention was turned to determine the capability of MONAL-41 to detect specific reductases in cell lysates. The A549 cell line was selected to investigate the applicability of MONAL-41 for the detection of reductase activity in cell extracts as it expresses CBR1 and several members of AKR superfamily, including AKR1A1, AKR1B10, AKR1B1, AKR1C3 [17, 24–27].

Figure 4 illustrates the dependence of the reductase activity profile in A549 cell extracts as a function of MONAL-41 concentration, clearly showing a decrease at substrate concentrations above 10 µM. This value, close to the concentration at which MONAL-41 starts inhibiting AKR1B10, together with the highest *kcat/KM* among the reductases tested, points out a possible prominent role of AKR1B10 in the MONAL-41-reductase activity of A549 cells which is in line with the expression profile of this AKR in A549 cells [17, 28]. A simple correlation between the enzyme activity rate of a cell extract and a specific enzymatic activity is challenging due to the overlapping substrate specificity of several reductases expressed in A549 cell line. However, we reasoned that performing the MONOL-41-assay in the presence of specific AKR inhibitors could allow separating their individual contribution to the overall reductase activity.

**Figure 4.**
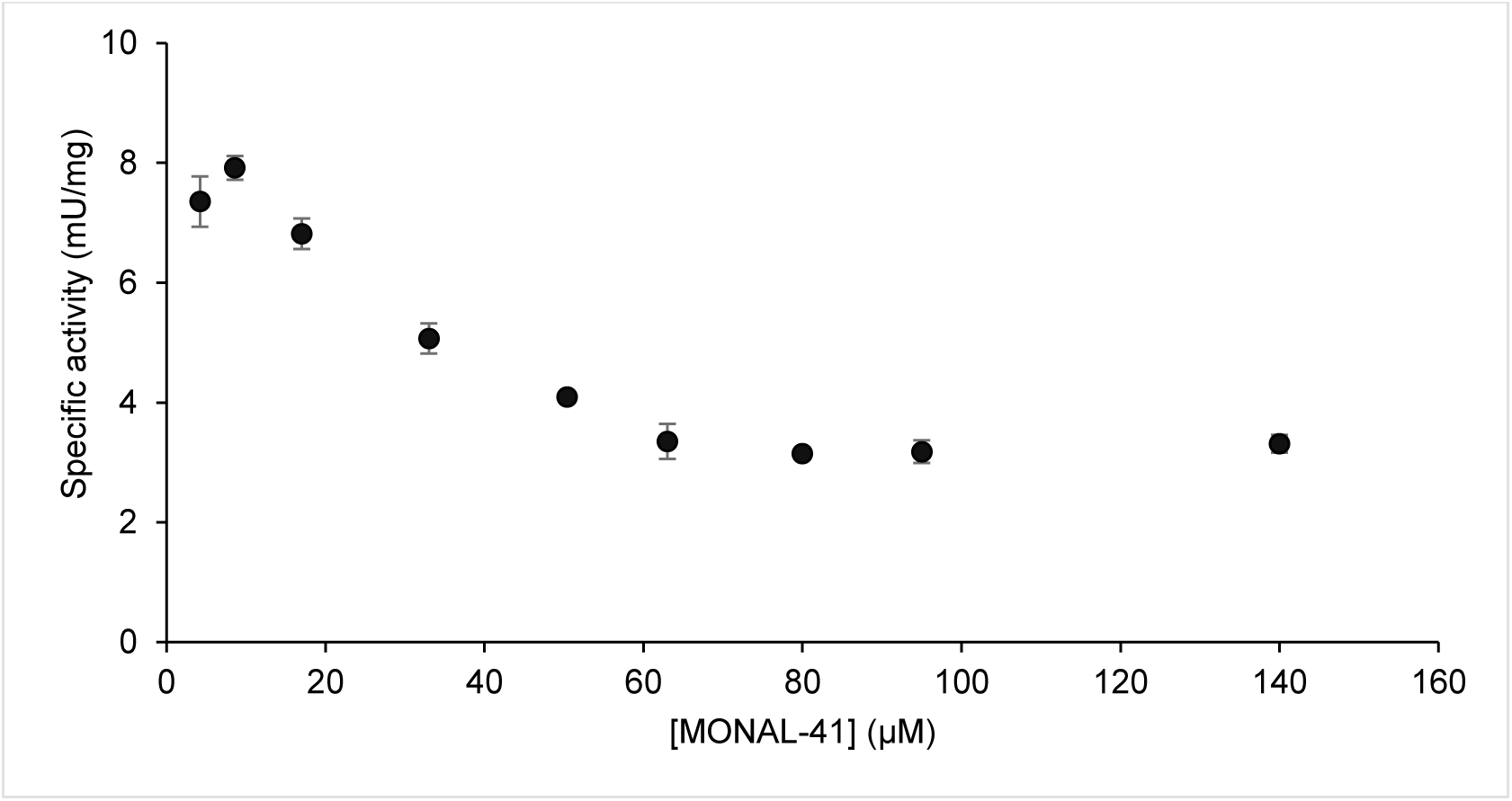
Dependence the enzyme activity rate of A549 cell extracts on MONAL-41 concentration. Results are reported as the mean ± standard deviation of three independent measurements.

To this end, cell extracts were treated with 3 µM tolrestat, a well-studied inhibitor of AKR1B1, AKR1B10 and AKR1A1 (IC50 values are 0.014 µM, 0.054 µM and 0.72 µM, respectively [29, 30]), which is ineffective against AKR1C3 and CBR1 [31]. When the experiment was performed in the presence of a low concentration of MONAL-41 (10.5 µM), a residual activity of 5.7 % with respect to the control was observed (Figure 5). Additionally, the assay was performed with high concentration (145 µM) of MONAL-41. Under these conditions, negligible activity of AKR1B10, AKR1A1, and AKR1C3 can be assumed due to substrate inhibition, as previously discussed. In these settings, where the only reductase that can be detected is CBR1, a residual activity of 7.9 % with respect to the control was observed (Figure 5).

**Figure 5.**
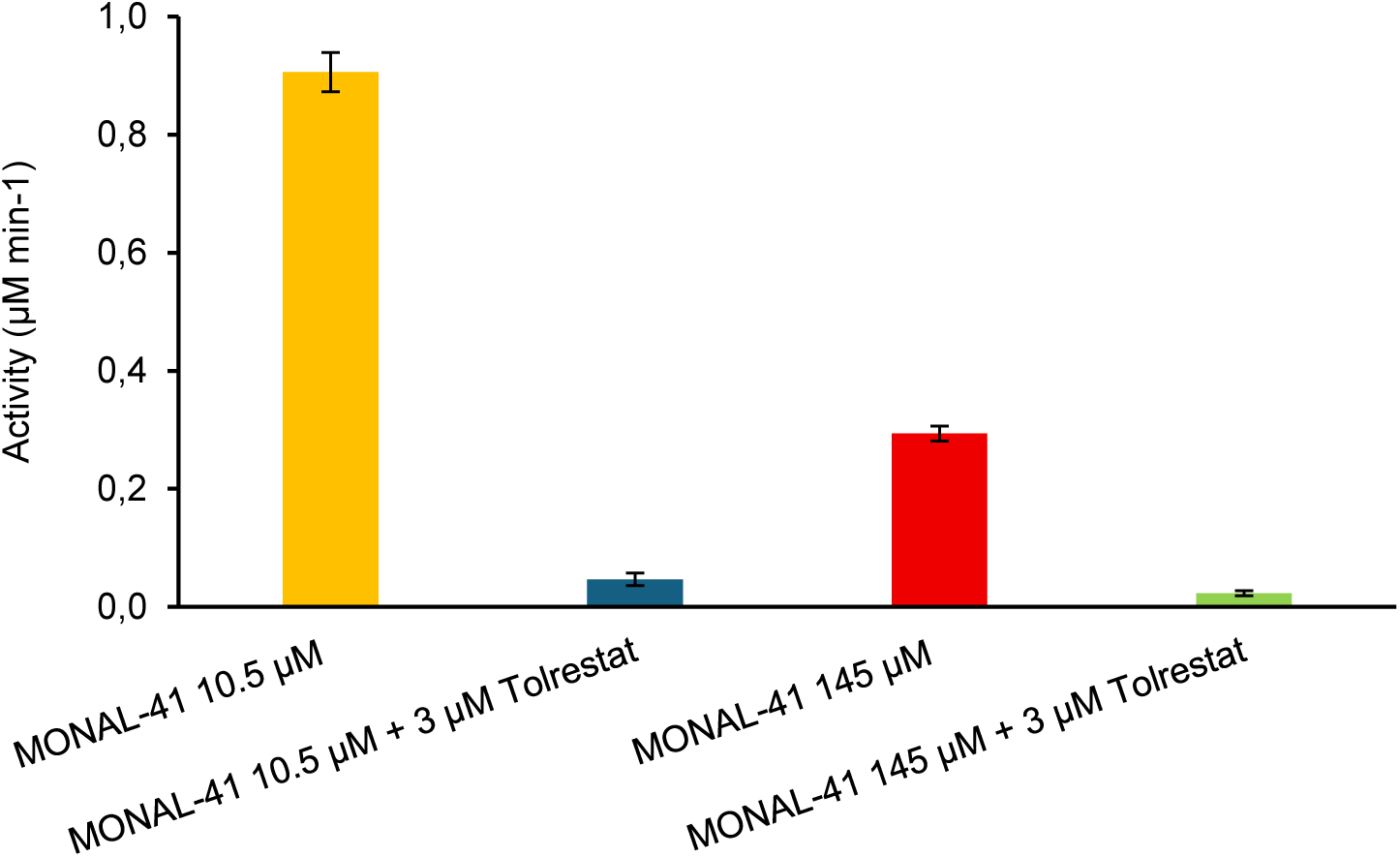
MONAL-41 reductase activity of A549 cell extracts. i. 10.5 µM MONAL-41 (yellow bar); ii. 10.5 µM MONAL-41 plus 3 µM tolrestat (blue bar); iii. 145 µM MONAL-41 (red bar); iv. 145 µM MONAL-41 plus 3 µM tolrestat (green bar). All the assay mixtures contained 0.7% (v/v) DMSO. Results are the mean ± standard deviation of three independent measurements.

This suggests that contribution of CBR1 to the MONAL-41-reductase activity of A549 cell extracts is reasonably low. A final consideration must be made based on the measured reductase activities in A549 cell extracts in the presence of 10.5 and 145 µM MONAL-41. As shown in Figure 5 and according to the activity profile of cell extract (Figure 4), reductase activity is lower at 145 µM compared with that observed at 10.5 µM. Comparison with the fitted experimental curves of purified enzymes (Figure 1) and their *kcat*/*KM* values suggests that AKR1B10, AKR1A1, and AKR1C3 can contribute to the MONAL-41 reductase activity at a substrate concentration of 10.5 µM in cell extracts, prior to the onset of substrate inhibition. AKR1B10 has a higher *kcat*/*KM* and expression in A549 cells [28] compared with AKR1C3 and AKR1A1. Concerning the possible contribution of AKR1C3, its markedly lower catalytic efficiency with respect to AKR1B10 makes it a poorly realistic player in MONAL-41 reductase activity in A549 cell line. On the other hand, even though the *kcat*/*KM* values of AKR1B10 and AKR1A1 are comparable, the reductase activity measured by MONAL-41 is likely correlated to AKR1B10 in this cell line, consistent with its higher transcript and protein abundance as previously reported [17, 28].

### 3.4. Detection of reductase activity in A549 living cells

Finally, we sought to investigate the possibility to use MONAL-41 assay for the detection of reductase activity in living cells. Preliminary experiments showed that MONAL-41 is not cytotoxic at concentrations up to 50 µM at 24 h. In the assay, A549 cells were treated with 50 µM MONAL-41, and the fluorescence emission of the conditioned medium of treated cells was evaluated at several time points ranging from 15 min to 4 h. The high membrane permeability of MONAL-41 and its product MONOL-41 (predicted XlogP3 = 3.1 [19] and XLogP3-AA: 2.3 [32], respectively) allowed direct evaluation in the medium. A clear increase in fluorescence emission was observed in comparison with a non-treated control, indicating that our approach is easily translatable to real-time measurements in living cells (Figure 6, panels A and B).

**Figure 6.**
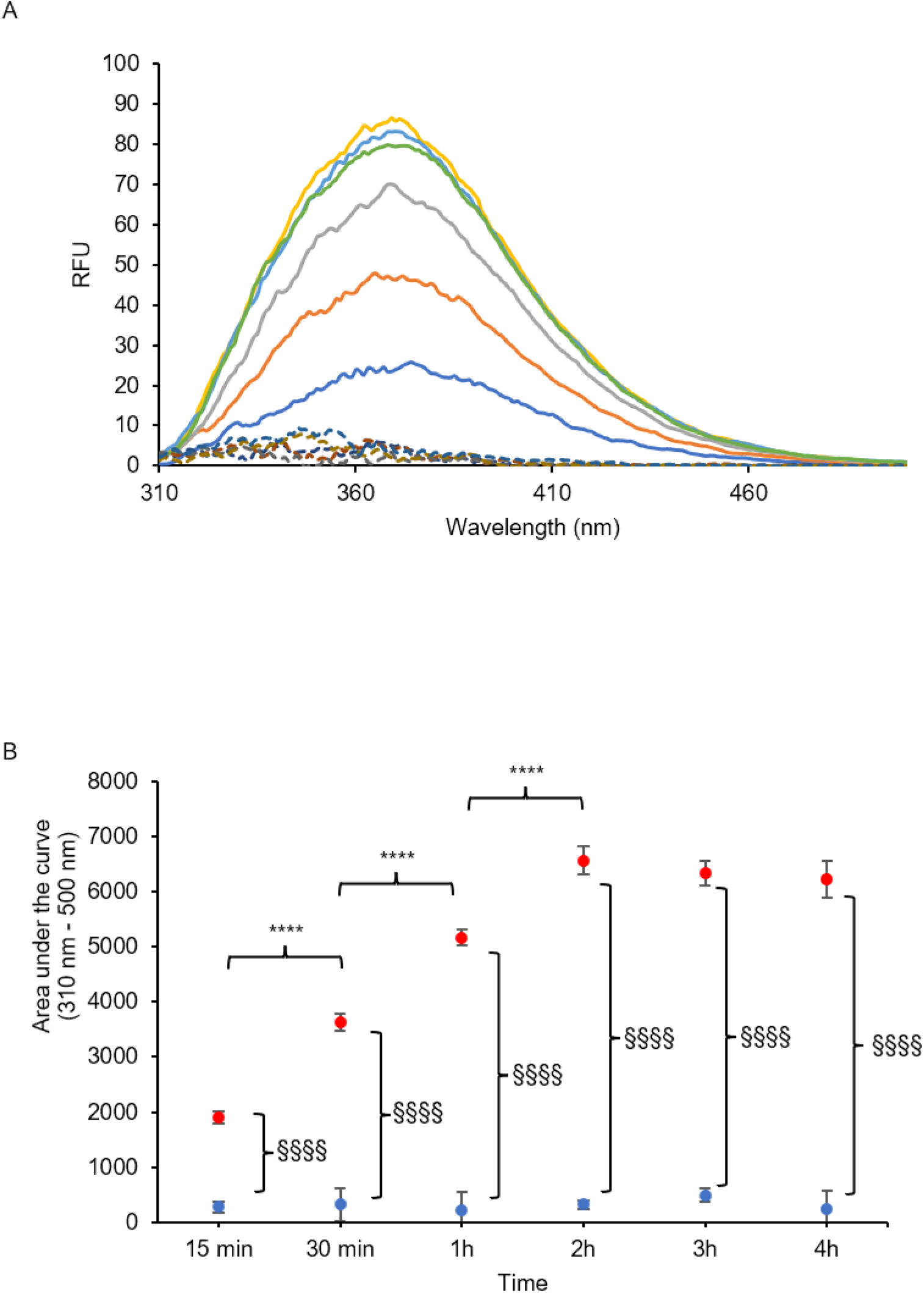
Detection of MONOL-41 in conditioned medium of A549 cell cultures by exciting at 296 nm. (A): Cells treated with 0.53 % (v/v) acetonitrile as controls (dashed lines) and cells treated with 50 μM MONAL-41(continuous lines) fluorescence emission spectra at 15 min (dark blue), 30 min (orange), 1 h (grey), 2 h (yellow), 3 h (light blue), 4 h (green). (B): Area under the curve of fluorescence emission peaks represented in panel (A), cells treated with 50 μM MONAL-41 (red dots) and cells treated with 0.53% (v/v) acetonitrile (blue dots). Error bars (when not visible, are within the symbols’ size) represent mean of three independent measurements (****= *p*<0.0001, refers to different time-points of treated cells; §§§§= *p*<0.0001, refers to difference between controls and treated cells at each time-point).

Compared with the more controlled conditions of dialyzed cell extracts, the whole-cell environment adds complexity due to the potential substrate oxidation mediated by aldehyde dehydrogenases, which are also expressed in the A549 cell line [33].

Depending on the relative abundance of reduced/oxidized cofactors and the expression of relevant enzymes, aldehydes may undergo either reduction or oxidation, potentially leading to the formation of MONOL-41 or its corresponding carboxylic acid, 4-methoxy-1-naphthoic acid (MONOIC-41).

Hence, we investigated the spectral properties of MONOIC-41, the oxidized form of MONAL-41 which was obtained by controlled oxidation with KMnO4[34]. Figure S2 reports the ESI-MS/MS spectral characterization of MONOIC-41. Absorbance and fluorescence spectra of MONOIC-41 (Figure S3) were largely superimposable with those of MONOL-41. The relative fluorescence quantum yield of the carboxylic acid form is approximately 8% compared to that of the target compound, and comparable with that of the MONAL substrate (Table 4).

**Table 4.** Relative Fluorescence emission *vs.* Absorbance ratios.

| Compound | Relative F/A % |
| --- | --- |
| MONOIC-41 | 8.45 |
| MONOL-41 | 100 |

Percentage calculated as $\frac{\int_{\lambda_{min}}^{\lambda_{max}} \text{Fluorescence}(\lambda) d\lambda}{\text{Absorbance}_{296 \text{ nm}}}$ relative to MONOL-41 value. Absorbance refers to the absorbance of the compounds at 296 nm obtained by a Biochrome Libra S60 UV-vis spectrophotometer (bandwidth = 1 nm; path length = 1 cm). Fluorescence spectra were obtained by exciting at 296 nm with a JASCO FT-750 fluorometer (PMT = medium intensity; bandwidth = 5 nm; path length = 1 cm).

This, combined with the ability of aldehyde dehydrogenases to oxidize MONAL-41 isomers [35] and their presence in A549 cells [33, 36], suggests further investigations of MONAL-41 as a substrate of aldehyde-oxidizing enzymes to better elucidate its use as an aldo-keto reductase biosensor in living cells.

## 4. Conclusions

In conclusion, we developed and optimized a microplate reader-based fluorometric assay which is more sensitive than the standard assay adopted for the detection of AKR1B10, and thus it could be useful to develop new inhibitor compounds. Additionally, the method is cost-effective and applicable for inhibitor screening. The assay is suitable for the detection of reductase activity in cell extracts and specific enzyme activities can be detected by combining the kinetic properties of the substrate with selective inhibitor treatments. This approach enables the deconvolution of individual enzyme activities in a cell extract through a few simple measurements.

## Abbreviations

AKR: aldo-keto reductase
AKR1B1: aldose reductase
AKR1B10: aldose reductase like
ALDH: aldehyde dehydrogenase
CBR1: carbonyl reductase 1
MONAL-41: 4-methoxy-1-naphthaldehyde
MONOL-41: 4-methoxy-1-naphthalenemethanol.

## CRediT authorship contribution statement

**Lucia Piazza:** Writing – original draft, Visualization, Methodology, Investigation, Data curation, Conceptualization, Formal analysis. **Raquel Pequerul:** Writing – review & editing, Methodology, Investigation, Formal analysis. **Xavier Parés:** Writing – review & editing, Supervision, Conceptualization. **Francesco Balestri:** Writing – review & editing, Validation. Supervision, Methodology, Funding acquisition, Conceptualization, Formal analysis. **Giovanni Signore:** Supervision, Methodology, Funding acquisition, Conceptualization, Formal analysis. **Antonella Del Corso:** Supervision, Methodology, Funding acquisition, Conceptualization, Formal analysis. **Jaume Farrés:** Writing – review & editing, Validation, Supervision, Methodology, Funding acquisition, Conceptualization, Formal analysis. All authors have read and agreed to the published version of the manuscript.

## Declaration of competing interests

The authors declare no conflicts of interest.

## Acknowledgements

This research was partially funded by European Union Next-GenerationEU-National Recovery and Resilience Plan (NRRP)-MISSION 4 COMPONENT 2, Spoke 6-CUP N. I53 C22000780001. This manuscript reflects only the authors’ views and opinions, neither the European Union nor the European Commission can be considered responsible for them. This research was also funded by the Spanish Ministerio de Ciencia, Innovación y Universidades (Agencia Estatal de Investigación, grant number PID2023-150696NB-I00 /MCIU/ AEI / 10.13039/501100011033 /FEDER, UE). RP obtained financial support from the company Advanced BioDesign through a research contract agreement with Universitat Autònoma de Barcelona.

## Appendix A. Supplementary data

Supplementary data to this article can be found online at

## Data availability

Data will be made available on request.

## Notes

### Competing Interest Statement

The authors have declared no competing interest.

