## Supplemental File for "A sensitive fluorometric assay to detect aldo-keto reductase and carbonyl reductase activity based on a naphthaldehyde derivative"

**Supplementary materials**

### S1. Optimization of assay to measure MONOL-41 production

The assays were performed in 96-well plates in a total volume of 100 µl/well. This approach is cost- and time-effective compared to *in-cuvette* procedures and minimize inner filter effects through both a reduced pathlength (3.3 mm) and top optics. As an additional precaution to avoid inner filter effects, the concentration of NADPH (0.08 mM) adopted in the assay mixtures of recombinant enzymes and cell extracts was reduced in comparison to the concentrations usually used in standard assay procedures (0.15-0.2 mM). Moreover, to minimize variations linked to the instrument sensitivity and improve the repeatability of measures, a DMSO solution of 1.8 µM MONOL-41 prepared in a separate well was used as an internal standard.

The spectral properties of MONOL-41 markedly differ from those of both MONAL-41 and NADPH, and the optimal excitation and emission wavelengths to maximize the signal of its emission were 296 nm and 370 nm, respectively [1]. However, the plate reader was equipped with absorption and emission filters (325 with a bandwidth of 40 nm and 410 with a bandwidth of 10 nm, respectively) that partially overlapped with the optimal set-up of excitation and emission wavelengths of both NADPH and MONAL-41, suggesting a potential interference in the detection of MONOL-41 emission. The quantum yield of MONAL-41 is negligible in aqueous media ($\Phi$=0.003 [1]), therefore a low, if any, impact on the fluorescence emission of MONOL-41 was expected. On the other hand, NADPH quantum yield is approximately 20 times lower [2] compared with that of MONOL-41 and it might impact on the signal of the targeted compound. The evaluation of the mutual influence of the two fluorophores on their respective signal emission showed that the variation of fluorescence linked to NADPH oxidation had a slight impact (approximately 1.3%) on the variation of fluorescence emission linked to the production of MONOL-41 (Figure S1).

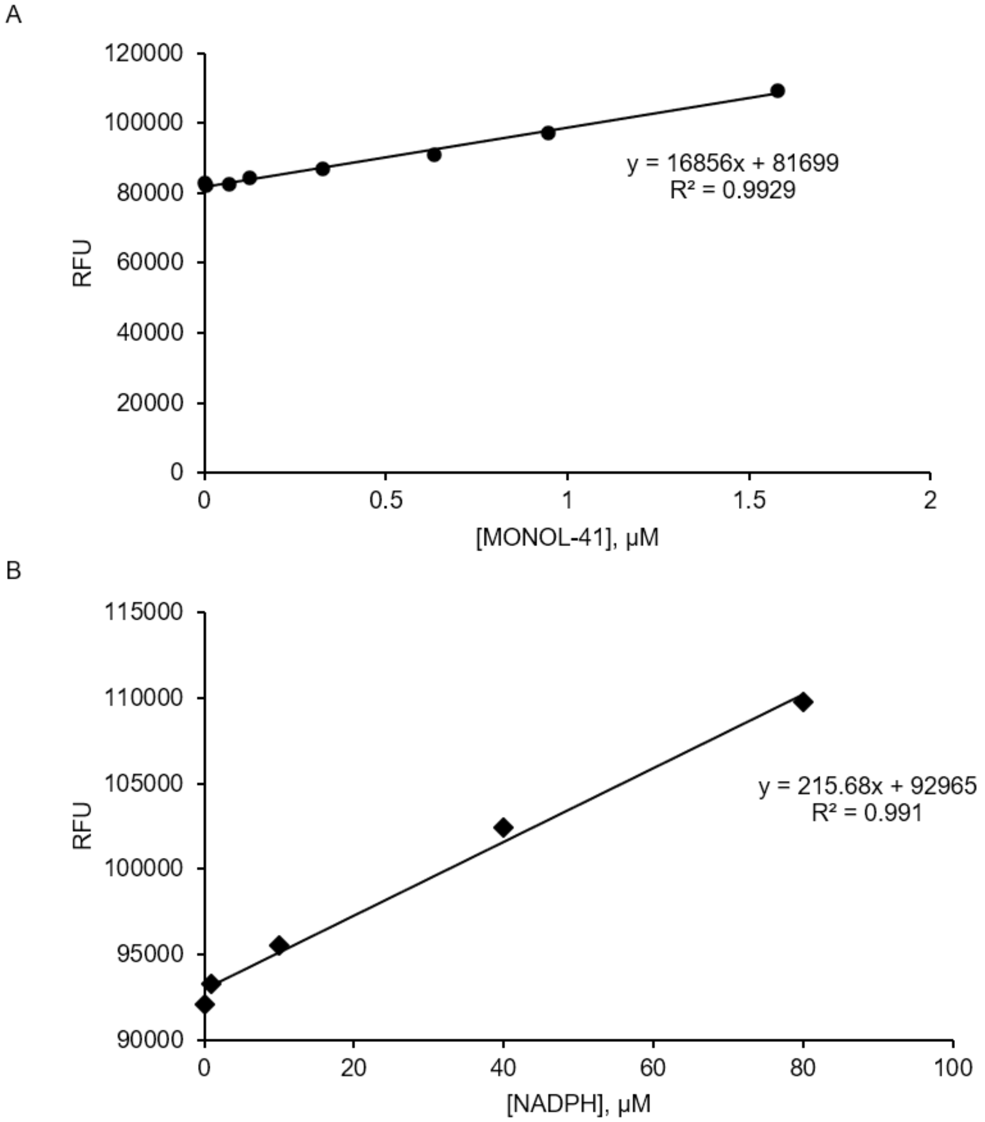


**Figure S1.** Calibration curve of MONOL-41 in the presence of 80 μM NADPH (panel A), and of NADPH in the presence of 1.58 μM MONOL-41 (panel B) obtained by exciting at 325-40 nm and detecting the emission of fluorescence at 410-10 nm (gain 1500).

### S2. Synthesis, identification and spectral characterization of MONOIC-41

MONOIC-41 was synthesized according to Shaabani et al. [3] with minor modifications. The oxidation of 1 mmol MONAL-41 in acetonitrile was obtained by adding 6.5 mmol of $KMnO_4$ and stirring the solution for 50 min. The solution was then centrifuged at 5000 *x g* for 8 min. The supernatant was collected and treated with a drop of hydrazine dihydrochloride to reduce any residual oxidant. The solution was centrifuged at 5000 *x g* for 8 min and the supernatant was collected. Five µL of the supernatant were resuspended in 495 µL of an acetonitrile/formic acid 0.1% (50/50, v/v) solution and analyzed by Orbitrap 120 Exploris mass spectrometer with ESI source in positive ion mode. Figure S2 reports the MS/MS fragmentation spectrum of the ion with m/z 203 corresponding to MONOIC-41. Fragments m/z 185 and m/z 159 corresponded to $[M\text{-}OH]^+$ and $[M\text{+}H\text{-}COO]^+$ and were in good agreement with typical fragmentation patterns of aromatic carboxylic acids. In addition, the resulting supernatant was diluted 1:280 in 100 mM sodium phosphate buffer, pH 7.0, and the absorption and emission spectra (Figure S3) were obtained through LIBRA S35 PC UV-visible spectrophotometer and JASCO FT-750 fluorometer (excitation wavelength= 296 nm; PMT = medium intensity; bandwidth = 5 nm), respectively.

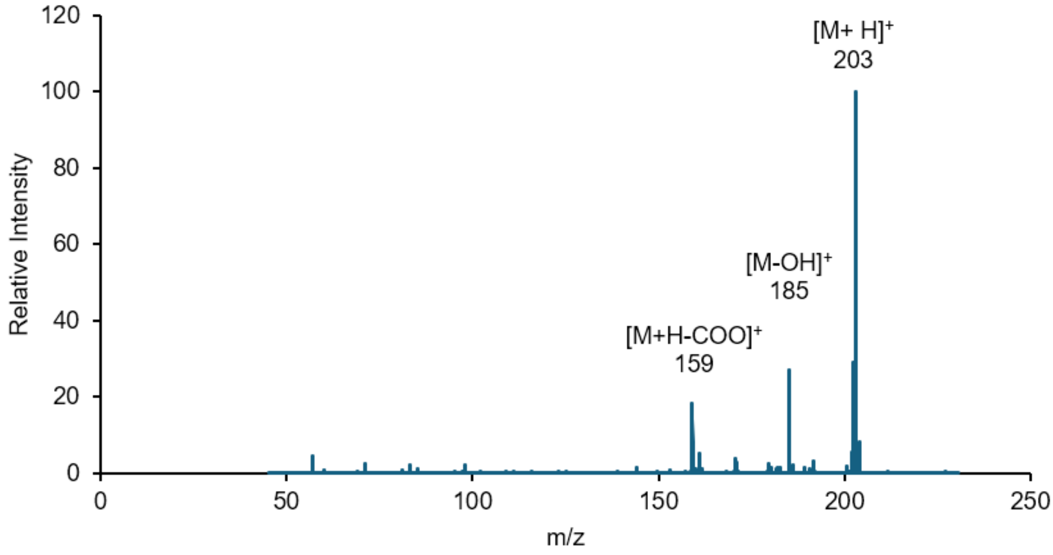


**Figure S2.** ESI-MS/MS fragmentation spectrum of putative MONOIC-41.

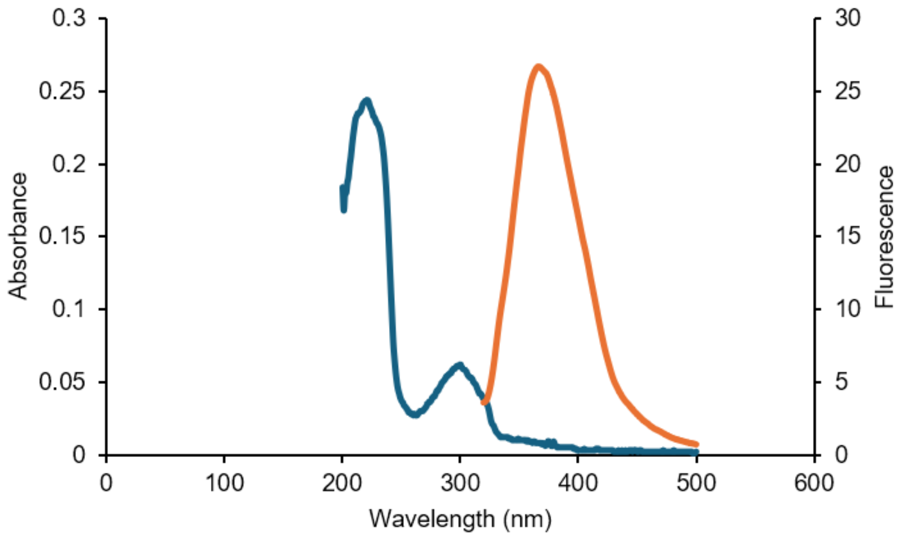


**Figure S3.** Absorption spectrum (blue line) and fluorescence spectrum (orange line) of putative MONOIC-41.

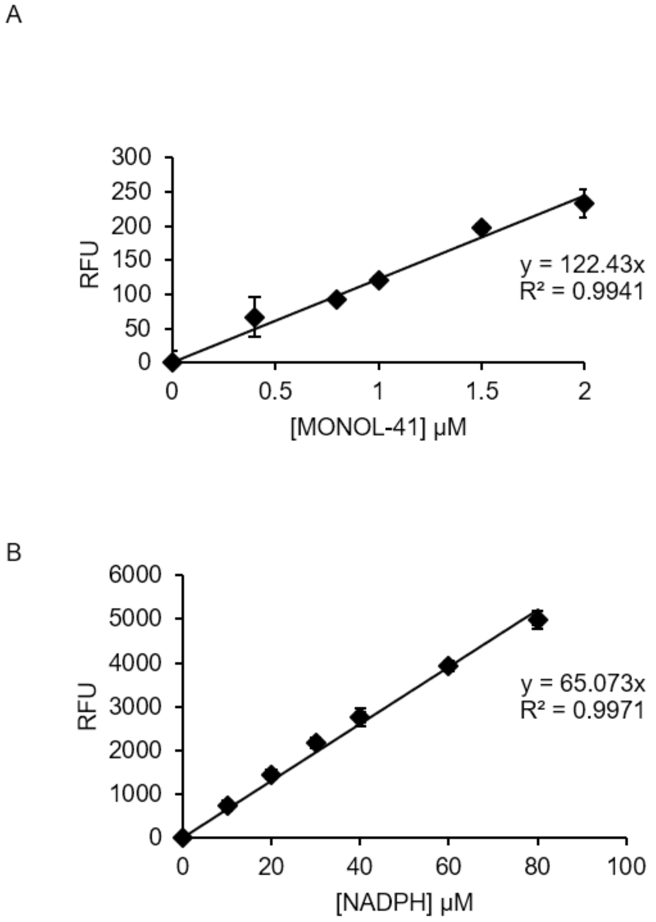


**Figure S4.** Panel A: Calibration curve of MONOL-41 in the presence of 80 µM NADPH, obtained by exciting at 320-10 nm and detecting the emission fluorescence at 410-10 nm (gain 1000); Panel B: Calibration curve of NADPH, obtained by exciting at 340-10 nm and detecting the emission fluorescence at 460 nm (gain 1000).
